# TM-Vec: template modeling vectors for fast homology detection and alignment

**DOI:** 10.1101/2022.07.25.501437

**Authors:** Tymor Hamamsy, James T. Morton, Daniel Berenberg, Nicholas Carriero, Vladimir Gligorijevic, Robert Blackwell, Charlie E. M. Strauss, Julia Koehler Leman, Kyunghyun Cho, Richard Bonneau

## Abstract

Exploiting sequence-structure-function relationships in molecular biology and computational modeling relies on detecting proteins with high sequence similarities. However, the most commonly used sequence alignment-based methods, such as BLAST, frequently fail on proteins with low sequence similarity to previously annotated proteins. We developed a deep learning method, TM-Vec, that uses sequence alignments to learn structural features that can then be used to search for structure-structure similarities in large sequence databases. We train TM-Vec to accurately predict TM-scores as a metric of structural similarity for pairs of structures directly from sequence pairs without the need for intermediate computation or solution of structures. For remote homologs (sequence similarity ≤ 10%) that are highly structurally similar (TM-score ? 0.6), we predict TM-scores within 0.026 of their value computed by TM-align. TM-Vec outperforms traditional sequence alignment methods and performs similar to structure-based alignment methods. TM-Vec was trained on the CATH and SwissModel structural databases and it has been tested on carefully curated structure-structure alignment databases that were designed specifically to test very remote homology detection methods. It scales sub-linearly for search against large protein databases and is well suited for discovering remotely homologous proteins.

## Introduction

A core tenet of molecular biology is the sequence-structure-function relationship that a protein’s amino-acid sequence determines its 3D structure which in turn determines its biological function. Detecting homology via sequence similarity is the standard approach to identifying evolutionarily conserved functions that are common between proteins [1, 2]. Over the last 50 years, sequence homology has enabled a wide array of applications, from annotating protein functions [3, 4, 5, 6, 7], predicting protein structure and protein interactions [8, 9, 10, 11, 12, 13], aiding protein design [14] and modeling evolutionary relationships [1].

Many standard sequence homology approaches are reliable for proteins that have high sequence similarity (>25%). However, unlike sequence homology, structural homology can be retained across long evolutionary time scales [15]. Recent estimates have suggested that over half of all proteins do not have detectable homologs in standard sequence databases due to their distant evolutionary relationships [16]. The challenge of remote homology detection is identifying these structurally similar proteins that don’t necessarily have high sequence similarity. It is widely understood that protein structure-structure alignments offer substantially more structure-function value at longer evolutionary distances that typically elude protein sequence alignment based methods. Using sequence-alignment based methods for closely related proteins and structure-alignment based methods for distantly related proteins could be an ideal hybrid approach that could offer substantially better sensitivity.

When protein structures are available, structural alignment tools such as TM-align [15], Dali [17], Fast [18], and Mammoth [19], can provide a measure of structural similarity by aligning protein structures via superposition [20, 15, 17, 19, 21]. While this approach can provide a measure of structural similarity in low-sequence similarity scenarios, there are two major limitations. First, protein structures are not available for most proteins. Considering rapid advances made by AlphaFold2, there remains a large gap between existing predicted structures (now around 200M) and the proteins that have been identified by the scientific community, with up to 68B proteins without known structures [22]. Furthermore, AlphaFold2 has limited utility in the context of predicting structures for novel folds [23] and for protein sequences with shallow multiple sequence alignments. Work on structure prediction that uses single or few homologous sequences is ongoing, but most methods exhibit reduced accuracy and take substantial time and memory resources per sequence that limits scaling to genomic protein databases.

Considering the rapid growth of protein structure databases, most existing structural alignment tools are far too computationally intensive to run at scale, requiring brute-force all-vs-all comparisons to query structurally similar proteins. While there are emerging tools for scalable homology search on structural databases [24], as well as for embedding proteins for either search or alignment [25, 26], no reliable tool exists that performs explicit structural-similarity search and alignment on large protein sequence databases.

To enable scalable structurally aware search on protein sequences, we propose a framework called TM-Vec, that can compute accurate structural similarity scores and structural alignments from pairs of sequences, while providing the ability to quickly index these results. Building upon recent advances in protein language models [27, 28, 29, 30, 31, 32, 33], we developed neural networks that can be fine-tuned on protein structures to (1) predict TM-scores between pairs of proteins using twin neural networks and (2) predict structural alignments between proteins using a differentiable Needleman-Wunsch algorithm. Our models output vector representations of proteins, which can be used to construct index-able databases to enable efficient querying of proteins by structural similarity.

We showcase the merits of our TM-Vec models in the context of CATH and SwissProt to show how our tool can scale with regards to database size while maintaining high precision in identifying structurally similar proteins. Our benchmarks suggest that TM-Vec can extrapolate beyond known fold space better than current state-of-the-art protein structure prediction systems. We contrast AlphaFold2 and TM-Vec in a case-study where TM-Vec can distinguish between bacteriocin classes more accurately than ColabFold in combination with TM-align [11]. TM-Vec is a broadly applicable tool that has the potential to enable the structural (and structure-similarity based) annotation of proteins and their functions in the vast and growing biodiversity contained in protein sequence collections.

## Results

Our contributions are two-fold: (1) we introduce a framework to perform scalable structure-aware search, TM-Vec, that affords substantial improvements in speed and sensitivity (Figure 1, Figure S1) [34] and (2) we introduce a differentiable sequence alignment algorithm, DeepBLAST, that performs structural alignments (Figure S2). The basis of our workflow is to predict the structural characteristics of proteins by training models on proteins with both sequences and structures available. Our alignment strategy uses recent developments in differentiable dynamic programming and protein language models to predict the structural alignments given by TM-align for pairs of protein sequences (Figure S2). For scalable search on large protein sequence databases, we introduce a separate twin neural network model that produces protein vectors that can be efficiently indexed and queried (Figure 1) [34, 35]. In order to encode structural information in these protein vectors, these twin neural networks are trained to approximate TM-scores (as a metric of structural similarity) of pairs of proteins. We showcase the ability of our workflow to extract structural information from protein sequences alone using a variety of benchmarks on the CATH, and SwissProt structure databases in addition to a case study on the BAGEL bacteriocin database.

**Figure 1:**
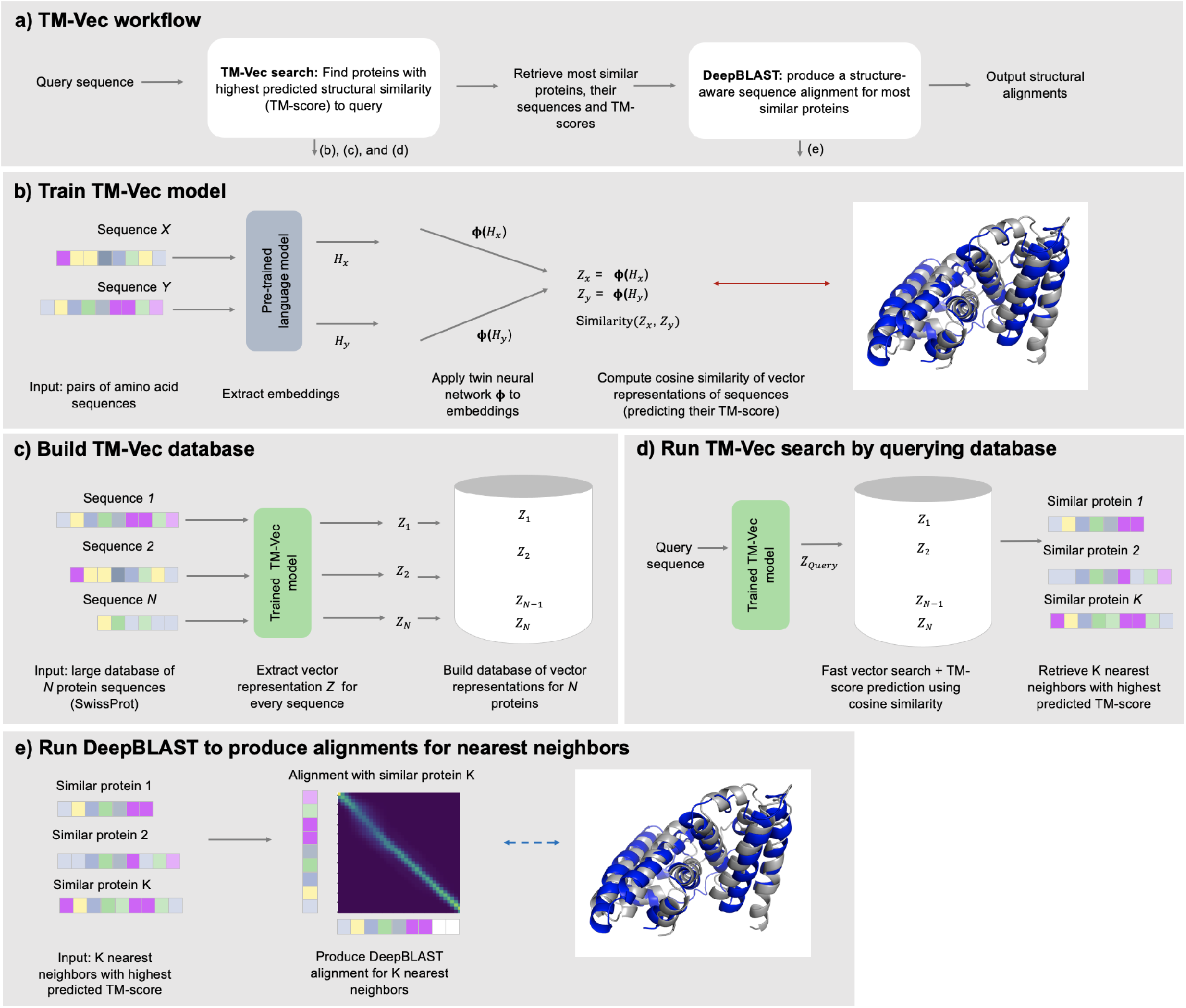
(a) The TM-Vec pipeline consists of two stages: retrieval and alignment. First, TM-Vec takes a query protein sequence and rapidly retrieves proteins that are predicted to have similar structures (TM-scores) to the query. Then, DeepBLAST produces alignments for the proteins with the highest predicted structural similarity. (b) TM-Vec is trained on pairs of amino acid sequences and their TM-scores. We first input a pair of sequences (i.e. domains, chains, proteins), and use a pretrained deep protein language model to extract embeddings for every residue of the sequence. Next, we apply a twin neural network, called *ϕ,* to the embeddings of each sequence, and produce a vector representation, *z,* for each sequence. The *ϕ* network is trained on millions of pairs of sequences and its architecture is detailed in Figure S1. Finally, we compute the cosine similarity of the vector representations, which is our prediction for the TM-score (structural similarity) of the pair. (c) We build a TM-Vec database by encoding large databases of protein sequences using a trained TM-Vec model. As an example, we input the sequences from SwissProt, extract vector representations for every sequence, and finally build an indexed database of TM-Vec’s structure-aware vector representations of proteins. (d) Demonstration of protein structure search using the TM-Vec pipeline. As the indexed database of vector representations has already been built, protein search consists of first encoding the query sequence using the trained TM-Vec model, and then performing fast vector search and TM-score prediction using cosine similarity as the search metric. As search results, we return the k-nearest neighbors with the highest predicted structural similarity (TM-score) with the query sequence. (e) As a last step, we apply DeepBLAST to produce structural alignments for the k-nearest neighbors to a query sequence.

### Extracting structural alignments from sequence using a differentiable alignment algorithm

We benchmarked DeepBLAST against three sequence alignment methods, Needleman-Wunsch, BLAST and HMMER in addition to four structural alignment methods that work directly with the atomic coordinates, FAST, TM-align, Dali and Mammoth-local (Table 1). TM-align achieves a global alignment by maximizing the 3D spatial overlap of the atoms in each protein. Conversely, the Mammoth local structure alignment scores feasible residue pairings between the proteins according to structural similarity of seven-contiguous-neighbor windows, as opposed to a remote homology philosophy where the full length structure is allowed to be flexible and does not require all the aligned atoms to overlap simultaneously after a rigid body orientation. Dali utilizes a distance matrix computed from hexapeptide contacts to align the two protein structures. FAST tries to preserve similar residue-residue contact patterns. We extracted the local structure alignment from the first phase of the Mammoth algorithm. These structure alignment algorithms span the range of expert opinions as to the most meaningful structure alignment (from emphasizing long-range overlap, contacts, and local-window similarity) and thus span potential disagreement across different prior approaches. No structure-alignment algorithms tested take sequence similarity into account.

**Table 1:** Malisam and Malidup Benchmarks. Sequence and structure alignment methods measured by their F1 score. Fast, TM-align, Dali Mammoth-local are structure-structure alignment methods and provide an structure-informed upper bound for this benchmark, as many of the most challenging alignments in this benchmark are ultimately structure-derived or curated with a structure-structure alignment as an oracle.

| Method | Malidup<br>F1 Score | # Detected | Malisam<br>F1 Score | # Detected |
| --- | --- | --- | --- | --- |
| BLAST | 0.019 ± 0.019 | 5 | 0.000 ± 0.000 | 2 |
| HMMER | 0.020 ± 0.02 | 8 | 0.020 ± 0.020 | 3 |
| Needleman-Wunsch | 0.098 ± 0.010 | 234 | 0.025 ± 0.003 | 129 |
| DeepBLAST | **0.144** ± 0.012 | 234 | **0.063** ± 0.007 | 129 |
| Mammoth-local | 0.483 ± 0.020 | 234 | 0.187 ± 0.017 | 129 |
| Fast | 0.569 ± 0.026 | 234 | 0.300 ± 0.030 | 129 |
| TM-align | 0.576 ± 0.024 | 234 | 0.393 ± 0.031 | 129 |
| Dali | **0.791** ± 0.014 | 234 | **0.619** ± 0.029 | 129 |

Our method DeepBLAST uses sequence alone; we do not supply the atomic coordinates of either protein to the algorithm after training it. To form a common reference for an optimal alignment, we focus on two gold standard benchmark sets comprised of manually curated structural alignments, named Malisam [36] and Malidup [37]. Manual structure alignment is intuitive human assessment typically emphasizing 3D overlap and topology preservation since those features are easier to visualize than a plethora of local alignments and contacts [38, 39, 40]. All methods tend to agree when the problem is trivial due to near sequence identity and near structural identity. Therefore the most valuable gold standard alignment benchmark set is where the dataset members have low sequence identity as well as varied degrees of structural similarity. Our benchmarks were performed on the curated Malisam [36] and Malidup [37] protein structural alignment benchmarking datasets (which are heavily skewed towards difficult-to-detect, low sequence identity remote homology).

As shown in Table 1, we observe that DeepBLAST outperforms all tested sequence alignment methods (Figure S3), but does not challenge the structural-alignment methods. In both benchmarks, the sequence similarity between proteins was below the observed detection limit for both BLAST and HMMER. As a result, these tools were not able to detect the vast majority of the alignments. This leaves Needleman-Wunsch as the baseline for sequence alignment methods. It is important to note again that there is no one definition of the best structural alignment [41, 42] and that this task becomes increasingly ambiguous as the remoteness of the homolog increases and the number of homologous residues declines. Thus the above F1 score tracks well with alignment accuracy but is limited in that it only scores sequence alignments with respect to a single reference alignment contained within the curated set.

### Scalable structural alignment search using twin neural networks

The challenge of applying our proposed structural alignment algorithm to large-scale protein databases is the demanding runtime requirements. Each TM-Vec structural alignment takes on the order of milliseconds and scales linearly with database size, making structural alignment searches on large databases impractical. To mitigate this issue, we developed TM-Vec, a model that is designed to efficiently query structurally similar proteins. Our strategy relies on the construction of twin neural networks, whose purpose is to provide per-protein vectors for fast indexing. The cosine distance of these vectors approximates the TM-score between pairs of proteins. This model can then be applied to entire protein databases to create an index over all of the protein vectors. The resulting database can be efficiently queried in *O*(log^2^ *n*) time for *n* proteins [34], providing sublinear scaling to retrieve structurally similar proteins based on their TM-score.

To evaluate the viability of our TM-score prediction strategy, we benchmarked TM-Vec on the SwissProt and CATH databases (Figure 2), and compared our approach to multiple state-of-the-art structure- and sequence-based methodologies. After training TM-Vec on approximately 150 million protein pairs from SwissProt (from 277K unique SwissProt chains), we observe a low prediction error (in the range of 0.025) that is independent of sequence identity across 1 million heldout protein pairs (Figure 2A). Like traditional sequence alignment methodologies, TM-Vec can accurately estimate structural differences when the sequence identity is greater than 90% (median error=0.005). Unlike traditional sequence alignment methods that typically cannot resolve sequence differences below 25% sequence identity [43], TM-Vec can resolve structural differences (and detect significant structural similarity) between sequence pairs with percent sequence identity less than 0.1 (median error=0.026). Overall, we see a strong correlation between the TM-scores predicted by TM-Vec and the TM-scores produced running TM-align (r=0.97, pval < 1 × 10^-5^) (Figure S4A).

**Figure 2:**
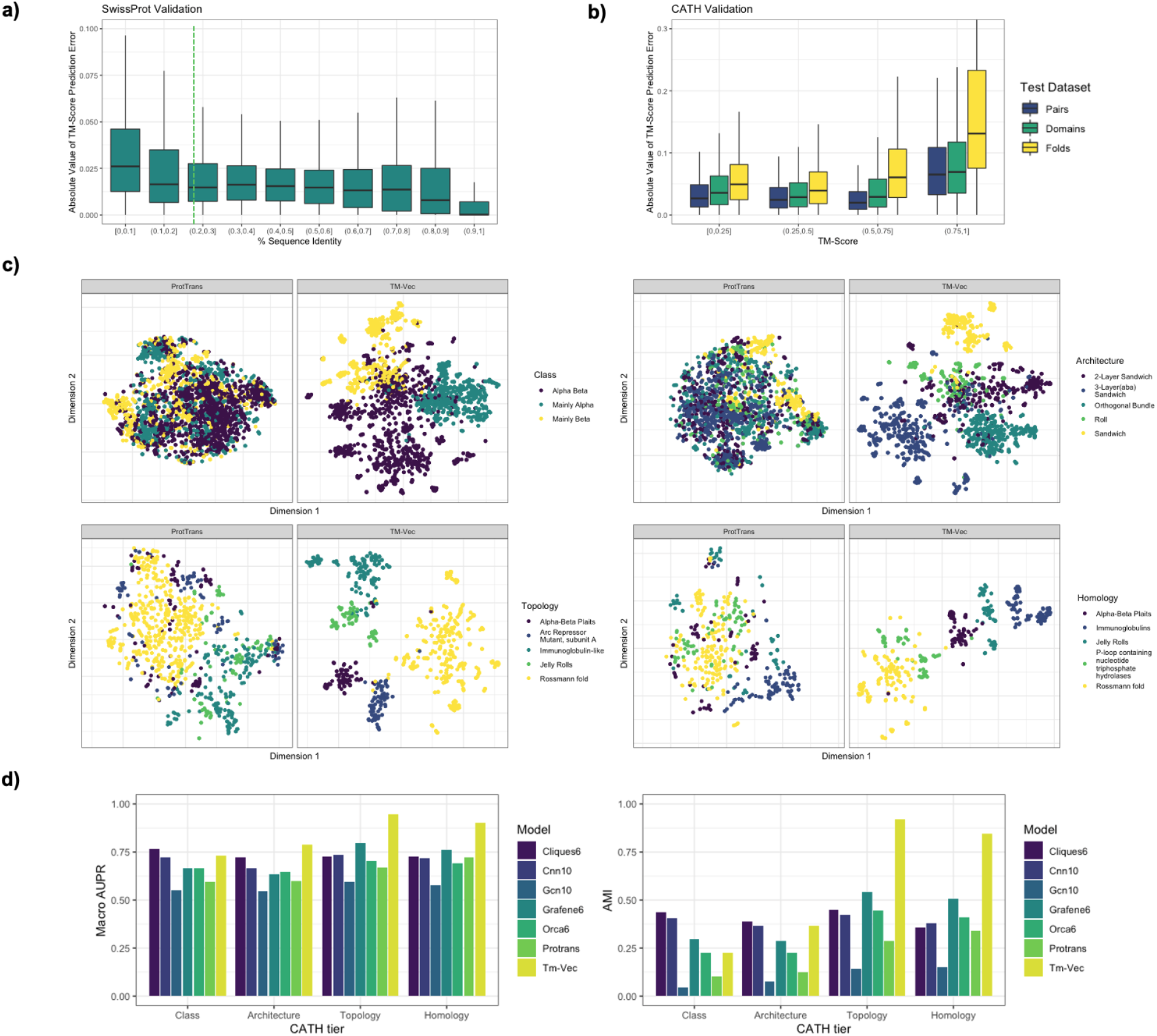
TM-Vec structural similarity prediction: Two TM-Vec models were built and benchmarked against protein sequence pairs from SwissProt and CATHS40. (a) SwissProt TM-score prediction errors (absolute value of difference between the known TM-score from running TM-align on structures and the TM-Vec predicted TM-score) for pairs with different sequence identities. Over 1 million pairs from a held-out test set are represented in this plot. Sequence similarity as measured by sequence identity ranges from [0, 0.1) (least similar) to (0.9, 1] (most similar). (b) TM-Vec absolute value of prediction error obtained from protein sequences compared to the TM-scores from TM-align obtained from protein structures. Prediction errors are stratified across 3 test benchmarking datasets: Pairs, Domains and Folds. The Pairs test dataset includes protein sequence pairs that were left out of model training/validation. Similarly, the Domains and Folds test dataset includes protein pairs derived from domains and folds that were never seen in model training/validation. (c) T-SNE visualization of protein embeddings from the top 5 most represented categories from each CATH classification tier (Class, Topology, Architecture, Homology) within the test dataset. For each CATH classification tier, TM-Vec embeddings are observed to separate structural categories better than the default protein sequence embeddings generated by ProtTrans. (d) Quantitative benchmarks of TM-Vec’s ability to predict CATH labels. We compare to ProtTrans, and 5 structure-based methods: cliques, GRAFENE, ORCA, DeepFRI, and GCN. Adjusted mutual information is computed from comparing spectral clustering assignments to structural label assignments for each CATH classification tier. Triplet scoring AUPR is a metric that determines how often cosine embedding distances from within structural categories are smaller than cosine embedding distances across structural categories.

We next validated TM-Vec on CATH protein domains that were clustered at 40% sequence similarity. For this, we validated TM-Vec’s predictions on three CATH held-out datasets: (1) pairs (of domains) that were never seen in training together; (2) domains that were both held out; and (3) folds that were both held out. We observe that TM-Vec accurately predicts the TM-scores for proteins from held-out pairs (r=0.936, pval < 1 × 10^-5^, median error=0.023) as well as held-out domains (r=0.901, pval < 1 × 10^-5^, median error=0.023) (Figure 2B, Figure S4B). TM-Vec’s prediction errors are highest for pairs with TM-scores in the [0.75-1.0] range, and its accuracy declines on held-out folds. However, the incremental increase in the generalization error for proteins in the held-out folds (r=0.781, pval < 1 × 10^-5^, median error=0.042) showcases that TM-Vec is robust to out-of-distribution observations, a critical requirement for extrapolating beyond the experimental structures within the PDB (Figure 2B, Figure S4B).

To further validate this finding, we applied TM-Vec to the Microbiome Immunity project (MIP) [44] that contains 200,000 *de novo* protein structure predictions from previously unknown proteins, including 148 putative novel folds. The correlation between our predictions and the TM-scores from MIP protein pairs with putative novel folds (r=0.785, pval < 1 × 10^-5^) was surprisingly close to the estimates we observed with the held-out CATH folds. Table S1 shows a confusion matrix for our TM-score predictions, where we observe that TM-Vec had a 99.9% true positive rate for predicting if a pair shared a novel fold (TM-score ≥ 0.5) and a false positive rate of 3.9%. Altogether, these validation benchmarks across SwissProt, CATH and MIP show that TM-Vec is well-suited to detect similarities between proteins with previously unknown protein structures and folds, extending the general utility of this work.

### Capturing structural information in the latent space

We visualize and benchmark the learned representations produced by TM-Vec against an array of alternative methods that depend on either sequence or structure alone. The results of our benchmarks show that TM-Vec implicitly learns representations that correlate well with structural classifications (Figure 2). Figure 2C demonstrates that TM-Vec embeddings capture the latent structural features of the CATH hierarchy. For comparison, embeddings produced by the pretrained language model that TM-Vec is based on, ProtTrans, are shown side by side with TM-Vec after training (Figure 2C). Across every tier of CATH, TM-Vec separates CATH structural classes more clearly than the default ProTrans embeddings.

To further evaluate the structural information of TM-Vec protein vectors, we encoded the CATH database using TM-Vec and performed search and classification. In our search benchmarks, we observe that TM-Vec is able to correctly retrieve proteins with the same fold in CATHS100 (97% accuracy) and CATHS40 (88.1% accuracy) for queried proteins (Table S2). In our classification benchmarks, we compared TM-Vec to several state-of-the-art methods (Cliques6, Cnn10, Gcn10, Grafene6, Orca6, and ProtTrans (see Online Methods)) using cluster adjusted mutual information and triplet scoring AUPR, to assess the representation quality of each method (Online Methods). We observe that TM-Vec outperforms the sequence-based and structure-based methods for Topology, Homology and Architecture classification as seen by its higher Macro AUPR for these tiers, indicating that TM-Vec is convolving both sequence and structure knowledge bases (Figure 2D). At the Class level, Clique6 and Cnn10 achieve higher Macro AUPR than TM-Vec. At the Topology level, TM-Vec had the highest Macro AUPR (Macro AUPR=0.94), and the second best method was Grafene6 (Macro AUPR = 0.79). The performance gap between the pretrained ProtTrans model (Macro AUPR = 0.66) and the fine-tuned model obtained with TM-Vec highlights the importance of fine-tuning with a structure-based objective. Furthermore, the fact that TM-vec outperforms sequence-based representations on the CATH dataset that is clustered at 40% sequence similarity provides evidence that TM-Vec learns quality structural features rather than a trivial feature of the underlying data or a function of sequence similarity.

### Remote homology detection and alignment: benchmarking on Malidup curated structure alignments

To gauge TM-Vec’s performance compared to existing structural alignment methods, we applied TM-Vec to the curated Malidup protein structural alignment benchmarking dataset [37], a difficult benchmark with low sequence identity and varied degrees of structural similarity. Each pair of proteins in this benchmark has a significant structurally similar region, a manually curated structure-structure alignment, and low sequence similarity that is either below or at the threshold of detection by sequence alignment tools. One of the challenges of benchmarking structural alignment methods is defining the ground truth structural alignment. As shown in Figure 3A, there are subtle disagreements between the manual alignments and the structural alignment methods, highlighting the uncertainty defining the optimal structural alignment. This is highlighted in scenarios where TM-align obtains a better structural superposition compared to the manual alignment (TM-align superimposes more atoms, or a greater extent of backbone regions than the manual alignment). All of the structure-aware methods agree at high structural similarity, TM-score=1 being perfect superposition of all atoms, but increasingly disagree as the TM-score declines. We observe that TM-Vec is directly comparable to structure-aware methods and the confidence bands for its trend line overlap with the trend lines for TM-align, and Fast (Figure 3A). Although the trend lines overlap, TM-Vec’s prediction errors have a higher variance than those of the structure-aware methods. To determine the agreement between sequence alignment methods and structural alignment methods, the TM-score was calculated for the predicted alignment. While DeepBLAST does not always generalize for divergent proteins, to illustrate an example where our method does obtain correct alignments for highly divergent proteins, we focused on two duplicated Annexin domains with a sequence identity of 24.7%. DeepBLAST accurately aligned these proteins (TM-score=0.75) and 4 of the 5 folds that were superimposed were in agreement with the manual alignment (Figure 3B-D). In contrast, Needleman-Wusnch was not able to identify any structural similarity between these two proteins (TM-score=0.33). The differences between Needleman-Wunsh and DeepBLAST are clear across all of the protein pairs in Malidup and Malisam. From the PSI scores shown in Figure S5, the high confidence alignments predicted by DeepBLAST are largely in agreement with the manually curated structural alignments. Furthermore, the sequence identity scores in Figure S5 reveal that DeepBLAST is able to obtain structural alignments for pairs that have ≤ 25% sequence identity, a known barrier for sequence alignment methods but can be resolved with the known protein structures. All together, these metrics suggest that DeepBLAST can perform local structural alignment.

**Figure 3:**
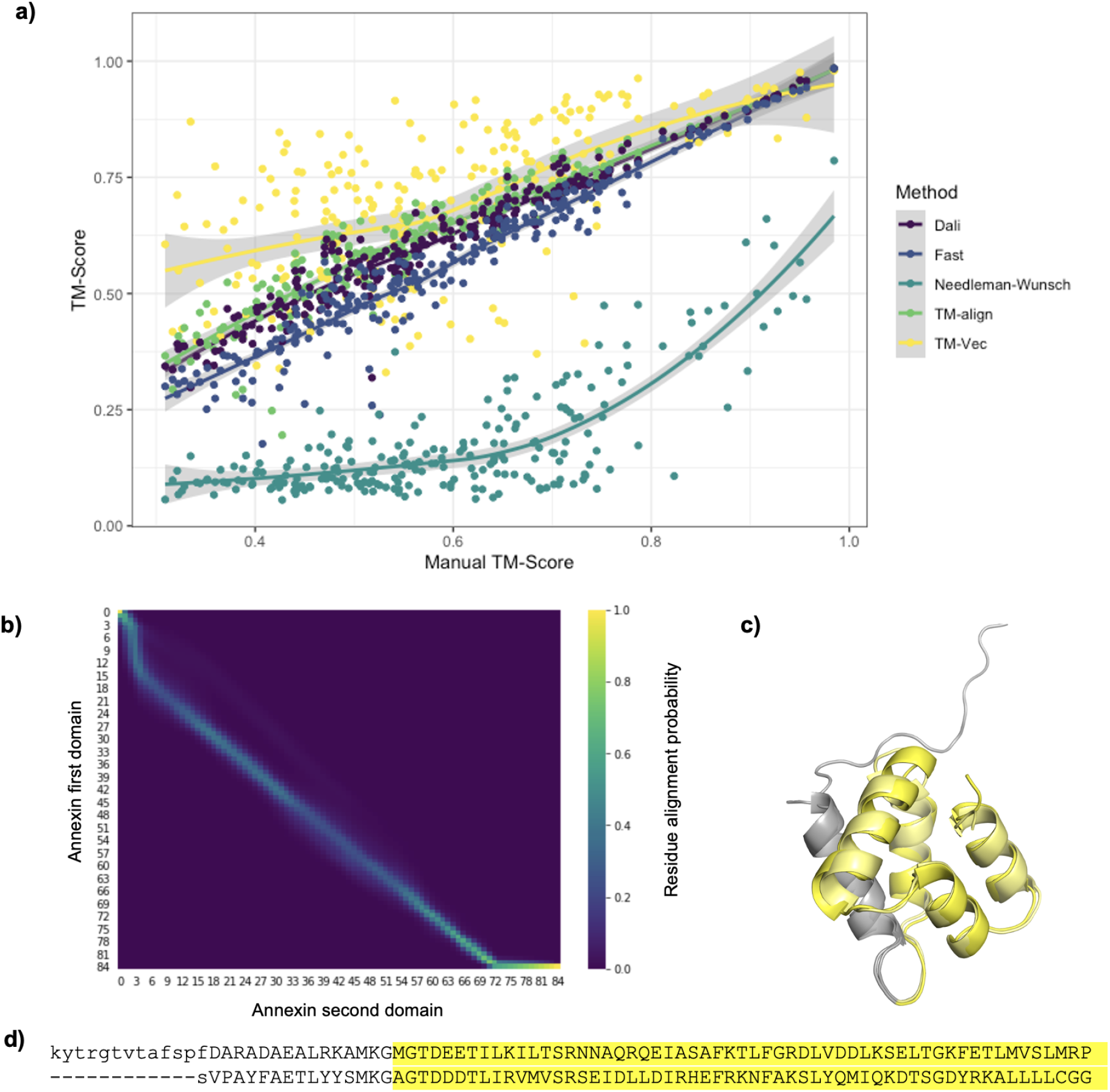
Annotating and aligning proteins in the Malidup benchmark. a) Comparison of multiple sequence and structural alignment methods. TM-Vec, and Needleman-Wunsch are both sequence alignment methods, whereas Fast, Dali, TM-align are structural alignment methods. Y-axis represents the predicted TM-score given a predicted alignment and the X-axis represents the TM-score from a manually curated alignment. TM-Vec performs comparable to structural alignment methods and outperforms Needleman-Wunsch. b) A predicted alignment of two duplicated Annexin domains from Malidup where DeepBLAST can accurately align (TM-score=0.75) and Needleman-Wunsch struggles with (TM-score=0.33). c) The manual alignment of the two duplicated Annexin domains with the agreement with DeepBLAST highlighted. Visualization of the manual structural alignment of the Malidup with the chains that DeepBLAST aligned correctly highlighted in yellow.

### Full repository level scaling and runtime

To show that TM-Vec can be applied to modern protein repositories, we benchmarked TM-Vec’s search runtime in multiple scenarios. After the creation of a TM-Vec database, a query is performed for a new protein sequence by first encoding it using the TM-Vec model, and then performing rapid vector search on the indexed protein TM-Vec database (Figure 1). Search runtimes for different numbers of queries and database sizes (Figure S6) empirically show that encoding queries is linear in time, with an ability to encode 50K queries on one GPU within 40 minutes (Figure S6A). In Figure S6B, we show sub-linear search performance. The search runtime benchmarks for different database sizes show that 50K queries on a database of 5M proteins can be performed within 20 seconds on a single GPU, showcasing that encoding sequences is the computational bottleneck in search. To constrast the TM-Vec query search time to existing sequence-based methods, we compared the TM-Vec query runtimes to Diamond [45] and BLAST. We observe that TM-Vec is not as fast as Diamond, which is optimized for short-reads and is known to have remote homology sensitivity and alignment performance similar to BLAST. TM-Vec does outperform BLAST in all cases, including in modes adapted for scaling TM-Vec described here, and our performance will scale sub-linearly with database size (Figure S6C). For example, TM-Vec achieves a 10X speedup compared to BLAST performing 1,000 queries on a database of 100,000 proteins, and this speedup will increase exponentially as the database size increases: on a 1M protein database there is a 100X speedup.

Thus, TM-Vec can be used to carry out full repository searches, large all vs all queries, and can do so with vastly improved remote homology detection and sensitivity. Further gains in computational performance are likely achievable (this work focuses on accuracy and sensitivity with respect to structure-structure quality alignments).

Once structurally similar proteins have been identified, structural alignments via the DeepBLAST can identify structurally homologous regions. Our structural alignment runtime benchmarks show that unlike the Needleman-Wunsch CPU implementations, the structural alignment runtime of our differentiable Needleman-Wunsch GPU implementation does not increase linearly with respect to the batch size, showcasing how our method can process multiple alignments in parallel on a single GPU (Figure S6D). Furthermore, we observe that both the CPU and GPU implementations scale linearly with regards to the length of both proteins, with our GPU implementation consistently yielding a 10X speed up over the CPU implementation.

### Case study: Bacteriocins

We conducted an analysis of a structurally diverse set of families, bacteriocins, using the BAGEL database [46]. Bacteriocins are peptide-derived molecules produced by bacteria, and often serve the role of antimicrobial peptides to target competing microbial species. They can also be involved in cell-cell communication. Several bacterial species encode bacteriocins and in light of their strong ecological benefits, bacteria are under evolutionary pressure to obfuscate these genes. As a result, bacteriocins showcase substantial sequence and structural diversity and are notoriously difficult to detect using sequence homology tools [47]. To date, less than 1004 bacteriocins have been identified and classified, despite there being trillions of microbial species [48] that have the potential to produce antimicrobial peptides.

Previous studies have shown that bacteriocin structures can be characterized by their highly modified polypeptides, suggesting structural cues to identify novel bacteriocins where sequence-similarity approaches fail. Our analysis revealed that TM-Vec can clearly partition bacteriocins according to their BAGEL annotations (Figure 4A, Figure 4A). Interestingly, unannotated bacteriocins identified in Morton et al [47] were identified to be structurally similar to Lanthipeptide A and B. In Figure 4C, we compared AlphaFold2 with TM-Vec on this bacteriocin dataset and found that TM-Vec distinguishes bacteriocin classes more accurately than ColabFold in combination with TM-align. Lastly, Figure 4D shows a DeepBLAST alignment for the 3 nearest classified bacteriocin neighbors of a putative bacteriocin identified by Hamid et al. [49].

**Figure 4:**
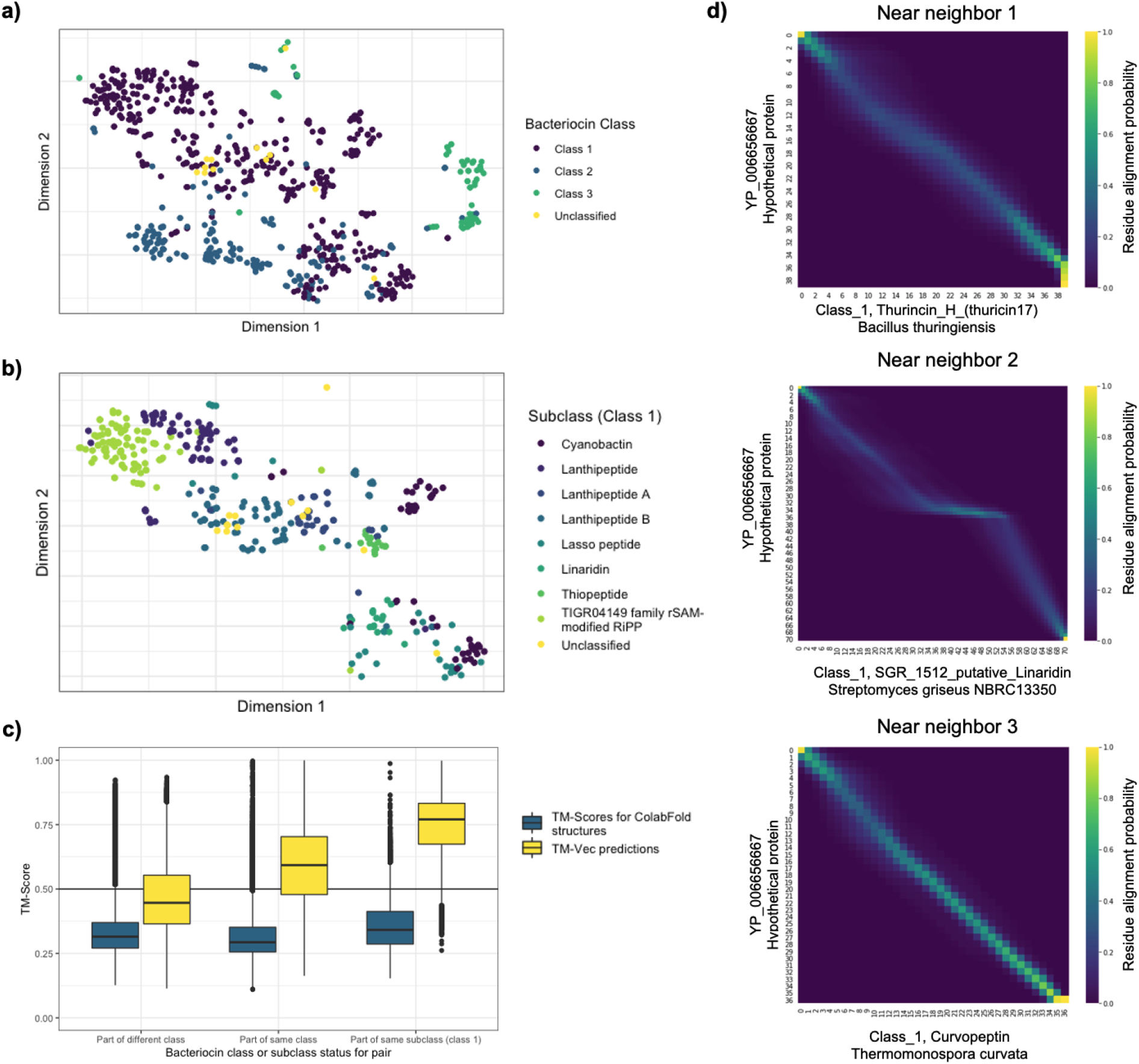
Annotating and aligning unclassified putative bacteriocins using TM-Vec. a) Visualizing TM-Vec embeddings using t-SNE for 3 classes of bacteriocins in addition to 28 unclassified putative bacteriocins. For 94% of the annotated bacteriocins, the nearest neighbor to a classified bacteriocin is in the same bacteriocin class. b) Visualization of Class 1 bacteriocins by subclass, highlighting how TM-Vec can recover multiple levels of manual annotation without protein structures. c) A comparison of TM-Vec’s TM-score predictions with the TM-scores produced by running TM-align on ColabFold predicted structures for every pair of bacteriocins. Using a TM-score of 0.5 as a structural similarity cutoff, TM-Vec is able to distinguish pairs of proteins that are in the same class versus different classes and in the same subclass for class 1 bacteriocins, while TM-align on ColabFold structures is not. d) DeepBLAST alignments for a putative bacteriocin, YP_006656667, and its 3 nearest neighbors in embedding space (i.e. they have the highest predicted TM-scores).

In Figure S7A, we found that Non-toxins clearly separated from the different bacteriocin class clusters based on their structural similarity. We further tested our ability to distinguish bacteriocins by training a k-nearest neighbor classifier for bacteriocin classes and Non-toxins; the overall precision and recall of these classifiers were 98% and 93% respectively (Figure S7B).

## Discussion

We have shown that TM-Vec has the potential to close the outstanding gap between protein sequence and structural information by enabling structural alignments from sequence information and remote homology search on repository scale protein sequence databases. From our benchmarks on SwissProt and CATH, TM-Vec can accurately predict the TM-score to quantify the structural similarities across wide-spread structural diversity, including remote homologs that fall below the 10% sequence identity threshold. Compared to sequence-based and structure-based methods, TM-Vec can competitively differentiate tiers of the CATH hierarchy, despite not being explicitly trained to classify CATH classes. Furthermore, TM-Vec is able to predict structural similarity close to existing structural similarity methods, while being able to query structurally similar proteins with both higher accuracy and lower runtimes than BLAST. TM-Vec search scales sub-linearly with respect to protein database size, and can handle millions of queries on tree-of-life scale databases per day on a single GPU machine. In addition to measuring structural similarity, TM-Vec can provide structural alignments that compare to existing structural alignment methods. From the Malidup benchmark, while we observe that there are certain remote homologs that our aligner misses, we consistently outperform sequence alignment methods. When we applied TM-Vec to the BAGEL database, we were able to accurately cluster bacteriocins based on both their class and subclass labels, a task that AlphaFold2 struggles with. We also were able to confidently annotate 28 putative bacteriocins by finding their nearest class or subclass clusters. Out insights hint at the potential to lower the barrier for natural product discovery.

In light of the promising advantages TM-Vec provides over the existing methodologies, there are a few limitations to consider. TM-Vec is not well suited to detect structural differences induced by point mutations. From a benchmark using the VIPUR dataset, TM-Vec was unable to detect structural differences caused by both deleterious and synonymous point mutations in proteins [50, 51, 52, 53, 54, 14]. Regarding structural alignments, DeepBLAST struggles to detect large insertions or deletions, which is commonly observed in remote homologs as suggested by TM-align generated training data. Recent advances incorporating linear affine gaps costs into differentiable dynamic programming algorithms [25] could play a role in resolving these challenges. Furthermore, integrating the TM-score prediction and the structural alignments into a multitask framework with a single pretrained protein language model may help boost the structural alignment accuracy.

Given the widespread biomedical applications and use cases of sequence search and alignment using tools such as BLAST, we anticipate that structural similarity search with TM-Vec will provide new opportunities for biological annotation. Due to the high structural precision and the fast query speed, TM-Vec and its future iterations are well poised to close the sequence-structure-function gap across the billions of observed proteins.

## Online Methods

We first describe our model that can produce structure-aware embeddings of protein sequences. We then use this model to build large searchable databases of protein representations that can be queried for finding proteins with similar structures, using only their sequence information. In the last piece of our pipeline, we produce protein structure alignments using sequences alone for the proteins that are predicted to have the most similar structures.

### TM-Vec Search

#### TM-Vec embedding model

The TM-Vec model is trained on pairs of protein sequences and their TM-scores (the measure of protein structure similarity we use), and leverages the latest advances in deep protein language models. When protein sequences are fed into the pipeline, a pretrained deep protein language model ProtTrans (‘ProtT5-XL-UniRef50’) is used to produce embeddings for every residue in the protein [32]. These residue embeddings are then fed into a twin neural network that we train, called *ϕ*. Figure S1 shows the function *ϕ* which takes residue embeddings and produces a flattened vector representation for each protein. *ϕ* is composed of several transformer encoder layers, followed by average pooling, dropout, and fully connected layers. Finally, we calculate the cosine distance between the reduced representations of each protein in the pair, and our training objective is to minimize the L1 distance between the cosine similarity of the pairs’ reduced representations, and their TM-score. Therefore, for any pair of protein sequences, a forward pass of our model can predict the pair’s TM-score, and can also be used to produce structure-aware embeddings for each protein sequence.

#### TM-Vec database creation

In order to build a large database of structure-aware protein embeddings, we start with large databases of protein sequences. Among the databases of protein sequences that we encode are SwissProt, and CATH. After encoding each protein sequence, we build an indexed vector-searchable database of protein embeddings using the FAISS package [34]. When this database is queried with a new sequence, we first embed the protein using a forward pass of the TM-Vec embedding model, and then we return the nearest neighbors of the query according to cosine similarity (the proteins in our database with the highest predicted structural similarity or TM-score). While one of our goals is to return the nearest neighbors in structure space for any query proteins, another goal is to also include the structural alignments for the nearest neighbors with the query protein, using sequences alone.

#### DeepBLAST alignment module

The DeepBLAST module uses a differentiable Needleman-Wunsch algorithm (Figure S2. Proteins *X* and *Y* are fed into the pretrained LSTM protein language model [27] to obtain embeddings *H_X_* and *H_Y_*. These residue-level embeddings are then propagated through the match embeddings (M) and gap embeddings (G) in order to obtain the match scores ***μ*** and the gap scores ***g***. The match and gap scores are used to evaluate the differentiable dynamic programming algorithm and generate a predicted alignment traceback. These alignments can then be fine-tuned using a training dataset of ground truth alignments.

##### Protein Language Modeling for alignment

In order to obtain an alignment from dynamic programming, the scoring parameters for matches and gaps must be obtained. Here we can use a number of pretrained protein language models to estimate these scoring parameters. These pretrained models ultimately construct a function, mapping a sequence of residues, represented as one-hot encodings, to a set of residue vectors, providing an alternative representation of these proteins. Often these models will learn these representations by being trained to predict randomly masked residues within a protein sequence. Multiple studies have shown the merits of these models when performing protein structure prediction, remote homology and protein design [29, 55, 28, 31, 30, 32, 33]. Here, we have used the pretrained LSTM PFam model from [27]. Using this pretrained language model, two proteins *X* and *Y* can be represented by embeddings ***H_X_*** ∈ ℝ^*p×d*^ and ***H_Y_*** ∈ ℝ^*q×d*^, where *p* and *q* represent the lengths of proteins *X* and *Y* and *d* is the embedding dimension of the language model. Given these representations, we can construct mappings *M* and *G* to obtain match scores and gap scores for the differentiable dynamic programming as follows

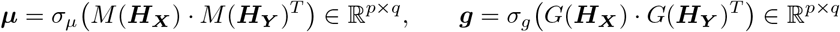

The functions *M*: ℝ^*t×d*^ → ℝ^*t×d*^ and *G*: ℝ^*t×d*^ → ℝ^*t×d*^ are intermediate functions that take in as input a set of *t* residue vectors. These functions are parameterized by LSTM networks, which can be fine-tuned through the backpropagation enabled by the differentiable dynamic programming. Activation functions *σ_μ_* and *σ_g_* are softplus and logsigmoid functions to ensure that the match scores ***μ*** are strictly positive and the gap scores ***g*** are strictly negative. These constraints are used to penalize gaps and reward matches. This also helps enforce identifiability of the model, which we have found to improve the accuracy of the model in practice.

##### Differentiable Dynamic Programming

Our proposed differential dynamic programming framework doesn’t learn any parameters; it is designed purely to enable backpropagation to fine-tune the scoring functions *M* and *G*. Differentiable dynamic programming has been extensively explored in the context of dynamic time warping [56, 57]. Koide et al [58] and Ofitserov et al [59] suggested that a differentiable Needleman-Wunsch alignment algorithm could be derived, but its implementation has remained elusive. Here, we provide the first GPU-accelerated implementation of the differentiable Needleman-Wunsch algorithm.

Previous work [57] has shown that backpropagation can be performed on dynamic programming algorithms by introducing smoothed maximum and argmax functions. Doing so will enable the computation of derivatives while providing a tight approximation to the optimal dynamic programming solution. The traditional Needleman-Wunsch algorithm can be defined with the following recursion

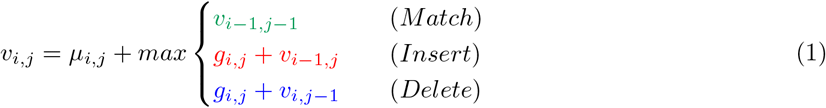

where the alignment score *v_i,j_* is evaluated on position *i* in the first sequence *X* and on position *j* in the second sequence *Y*. Sequences *X* and *Y* are of lengths *n* and *m* respectively. *μ_i,j_* represents the log-odds score of residues *X_i_* and *Y_j_* being aligned and *g_ij_* represents the log-odds score of an insertion or a deletion at positions *i* and *j*. Due to the structure of dynamic programming problems, *v_n,m_* is guaranteed to be the optimal alignment score between the two sequences. Furthermore, the optimal alignment can be obtained by tracing the highest-scoring path through the alignment matrix via argmax operations.

As neither the max nor the argmax operations are differentiable, the alignment scores and the traceback cannot be differentiated in the traditional formulation of the traceback operations needed to generate alignments. Accordingly, Mensch et al [57] introduced smoothed differentiable operators

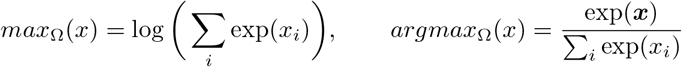

where the smooth max operator *max_Ω_*(*x*) is given by the log sum exp function and the smoothed *argmax_Ω_*(*x*) is given by the softmax function. Since the softmax function can be derived from the derivative of *max_Ω_*, the traceback matrix can also obtained by differentiating the resulting alignment matrix. The resulting traceback matrix will yield the expected alignment between the two proteins.

Since the loss function is defined as the difference between the predicted traceback matrix and the ground truth traceback matrix, the derivatives of the traceback matrix also need to be defined, which requires both the computations of the directional derivatives and the local Hessians of the alignment matrix (Algorithm 2).

###### Algorithm 1 Compute DeepBLAST_Ω_(*θ*) and ∇DeepBLAST_Ω_(*θ*)

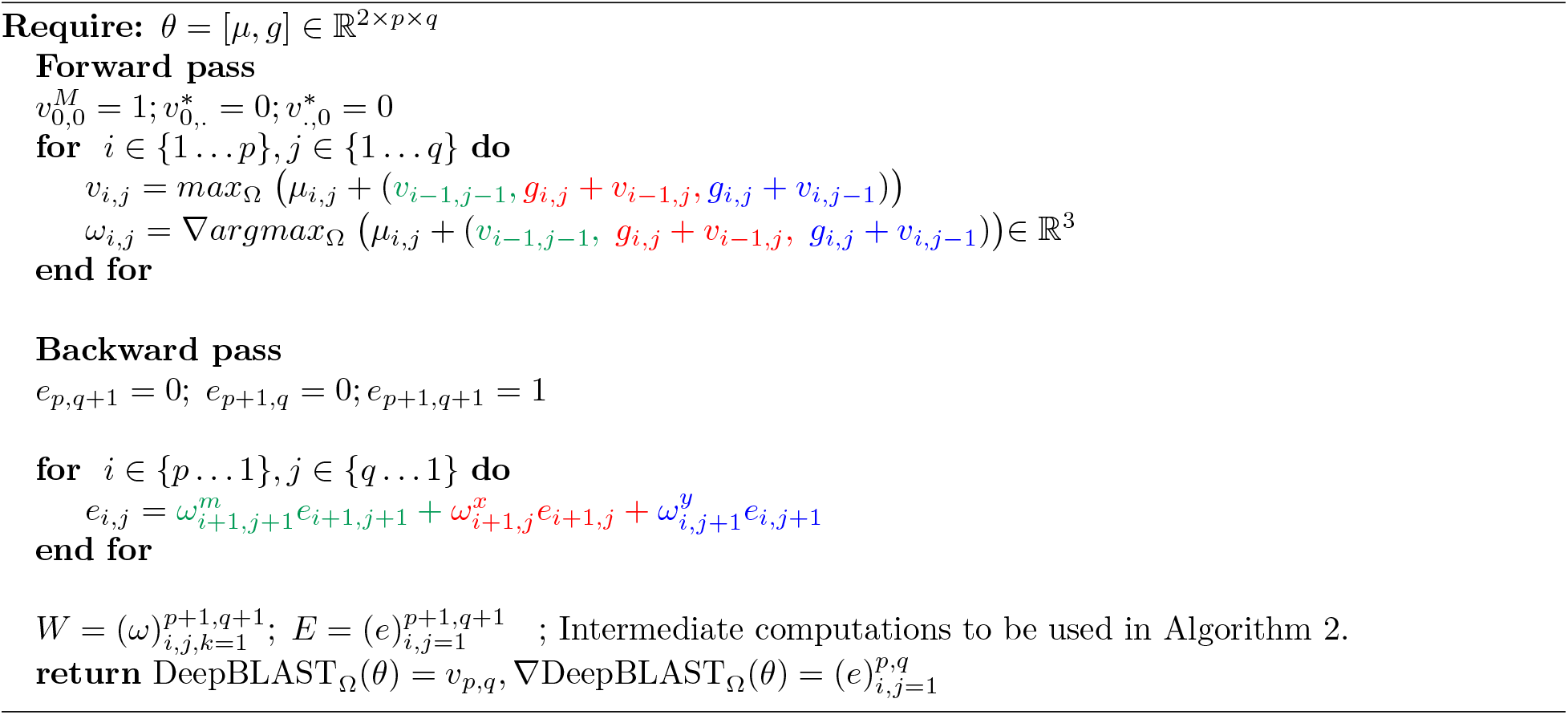

###### Algorithm 2 Compute 〈∇DeepBLAST_Ω_(*θ*), *Z*〉 and ∇^2^DeepBLAST_Ω_(*θ*)*Z*

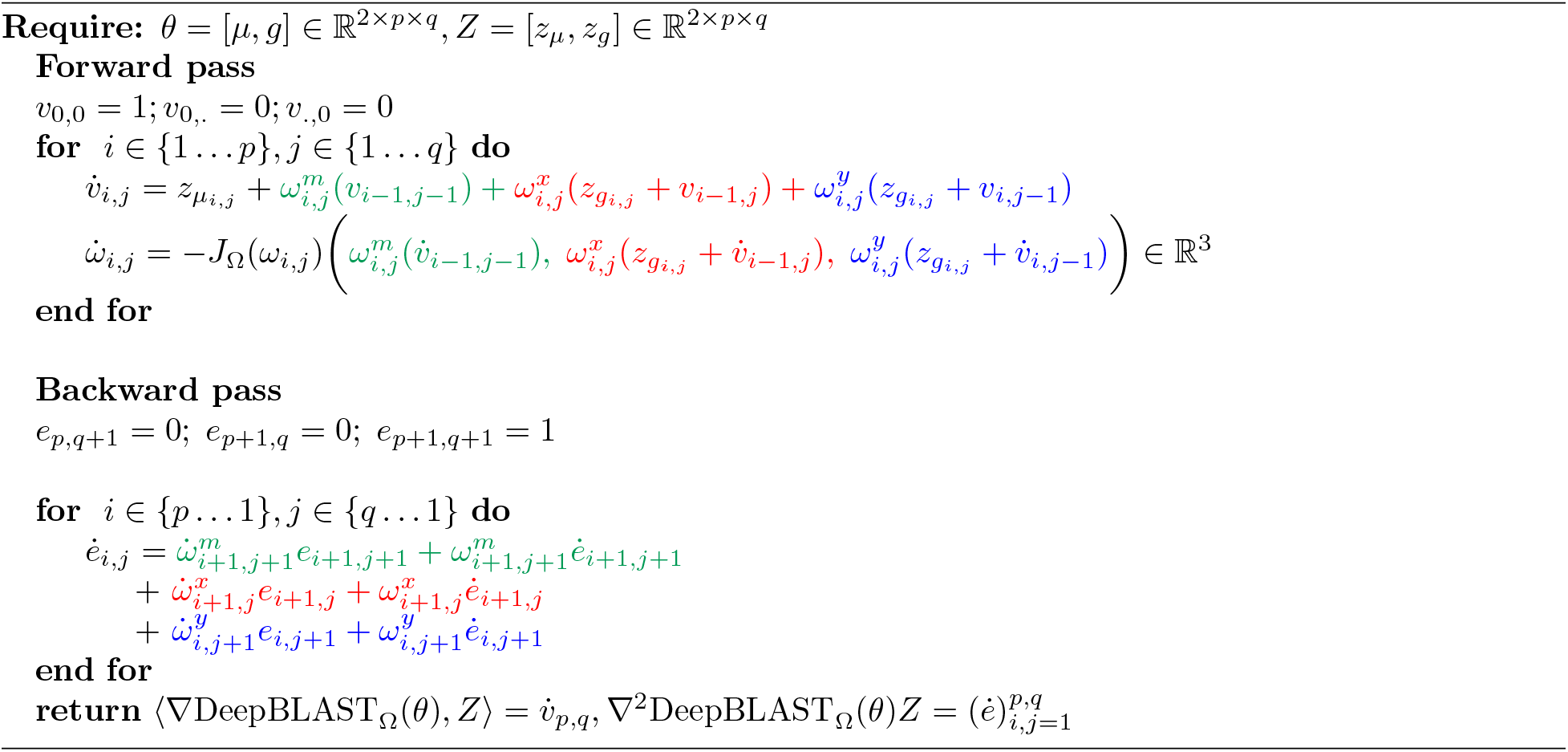

In practice, dynamic programming can be the major computational bottleneck due to GPU data transfer and the quadratic runtime of the Needleman-Wunsch algorithm. To address this, we have implemented a GPU-accelerated differentiable Needleman-Wunsch algorithm inspired by Manavski et al [60]. As can be seen from the benchmarks shown in Figure S6D, this algorithm is an order of magnitude faster than the naive CPU-bound Needleman-Wunsch implementation. Furthermore, this algorithm can enable batching, allowing for multiple alignments to be processed in parallel. As shown in Figure S6D, larger batch sizes can further improve the scaling over CPU-bound alignments.

##### Alignment Loss Function

By defining a loss function between the predicted alignment and the structural alignment from TM-align, we can evaluate the accuracy of DeepBLAST and fine-tune the functions *M* and *G*. Mensch et al [57] proposed using the Euclidean distance between the predicted and ground truth alignments as a loss function. In practice, we found that a cross-entropy loss provided more reasonable alignment results. This loss is given by

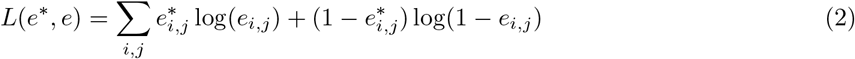

where *e*\* is the ground truth alignment and e is the predicted alignment. As shown in [57], the predicted traceback matrix represents the expectation across all possible predicted alignments, which is represented as a matrix of probabilities. As a result, the resulting alignment problem can be interpreted as a classification task of identifying whether two residues between a pair of proteins are alignable. This provides additional motivation for utilizing cross-entropy as a loss function.

#### Datasets

##### TM-Vec Search

TM-Vec is trained on pairs of protein/domain sequences along with data about the structural alignment for the pair. For every pair of proteins in our training dataset, we run the method TM-align, which is an algorithm for protein structure comparisons that is independent of protein sequences. TM-align produces a TM-score between 0 and 1, where a score below 0.2 represents a pair of unrelated proteins, a score above 0.5 implies that proteins are in the same fold, and 1 is a perfect match, indicating the same protein structure. Part of our pipeline involves validating whether our model can predict the TM-scores of pairs of proteins.

##### Protein chain pairs dataset

The model that we ultimately use to encode protein sequences, is trained on pairs of protein chains. We sample pairs of chains from SwissModel, which has over 500K chains in it. We filter out protein chains that are longer than 300 residues, and are left with 277K chains. With these chains we make pairs, we also make sure to oversample pairs of proteins with similar folds, which we do by using information from Gene3D about the predicted domains within protein chains. The dataset that we end up with has 150 million pairs of protein chains. We run TM align for each one of these pairs of protein chains by using their structures from the PDB. We pull out the TM-scores and sequence identity for every pair of chains. Lastly, we split our dataset in train/validation and test splits. Our held out test dataset has 1 million pairs, and our train/validation split (which we randomly split into training/validation during our training) has 141 million pairs.

##### Domain pairs dataset

In order to validate whether our model can approximate TM-scores for domains and remote homologs, we build a dataset of pairs from the heavily curated CATH domains dataset. We start with the CATH non-redundant dataset of protein domains with no more than 40% sequence similarity. This dataset comprises 31K protein domains. We then sample for domains that are filter out domains that are longer than 300 residues, which leaves us with 30K domains. While all pairwise-combinations of these 30K domains would lead 450 million pairs, we aim to build a balanced dataset, and dissimilar protein structures represent the vast majority of pairs (i.e. domains with very different folds). Therefore, we undersample pairs of CATH domains that come from different folds. The CATH dataset that we use for our various experiments includes 23 million pairs of domains.

We further split this dataset into training/validation and testing splits, and we evaluate our performance on CATH S40 on left out domain pairs (in the domain pair, one domains is in training/validation, while the other domain is in the held-out dataset), left out domains (both domains are not in the training/validation dataset), and left out folds (both domains come from folds that are not in the training/validation dataset). Our training/validation dataset contains 19 million pairs; our left out pairs dataset contains 100k pairs, our left out domains dataset contains 100K pairs; and our left out folds dataset contains 500K pairs.

##### Malidup and Malisam datasets

Part of our sequence alignment benchmarks were performed on the curated Malisam [36] and Malidup [37] protein structural alignment benchmarking datasets. We also used Malidup to benchmark TM-Vec and DeepBLAST. Malidup consists of 241 pairwise structure alignments for homologous domains within the same chain. These pairs are structural similar remote homologs. Malisam consists of 130 pairs of analogous motifs.

##### Structure alignment dataset

We trained DeepBLAST on 1.5M alignments from the PDB [61] obtained using TM-align [62]. These proteins were obtained from a curated collection of 40k protein structures [63]. Details behind the model specification and training can be found in [64].

##### Bacteriocins dataset

The bacteriocin sequences and metadata we use come from the bacteriocin database, BAGEL [46], and the putative unannotated bacteriocins that we use come from Morton et al [47].

##### Microbiome Immunity Project (MIP) novel fold dataset

In this project, there were protein structure predictions for 200,000 diverse microbial protein sequences, representing 161 putative novel folds, and the authors calculated TM-scores for pairs of proteins with novel folds [44].

##### Embedding methods data

The data we used for this evaluation is CATH NR-S40 dataset (NR-S40) [65]- a collection of approximately 30 thousand proteins of maximally 40% sequence identity representing a diverse sampling of each tier in the CATH hierarchy. The dataset is partitioned into train, validation, and test sets. All of the benchmarks are conducted on the test set and all trainable methods in the comparison study were trained using the train and validation sets.

#### TM-Vec training

TM-Vec has 17.3 million parameters, and is 199MB. We trained TM-Vec on 8 Nvidia V100 GPUs for 5 days. This represented 5 epochs of training.

### DeepBLAST training

The final DeepBLAST model consisted of 4 LSTM layers of dimension 512 to parameterize the match embeddings *M* and gap embeddings *G*. A 2 layer bidirectional LSTM protein language model pretrained by [27] was used as a precursor step for estimating residue vectors. The resulting model had a total of 100M parameters. We used the ADAM optimizer to train the weights with an initial learning rate of 5 × 10^-5^ and the pretrained LSTM model weights were frozen. A batch size of 160 alignments was used for training. DeepBLAST was trained for 10 epochs on 4 Nvidia V100 GPUs for 4 days.

### DeepBLAST alignment accuracy assessment

Alignment accuracy was assessed on a held out test dataset of 79k structural alignments. To determine how well TM-Vec generalizes, proteins that were in both the heldout LSTM PFam dataset [66] and the held out TM-align alignments used to train TM-Vec were analyzed. Within the TM-Vec held out dataset, 57,444 alignments were constructed from proteins that were unique to the TM-Vec held out dataset, 18,53 alignments contained proteins that were similar to those trained from the LSTM PFam training dataset and 19,967 alignments contained a single protein that was unique to the TM-Vec held out dataset and a single protein that was in the LSTM PFam training dataset. To evaluate the accuracy of the alignments, precision and recall were computed from the number of correctly identified matching residues. Since each alignment can be represented as a bipartite graph where the edges represents matching residues between two proteins, precision and recall can be extracted from comparing the edge sets between the predicted alignment and the known alignments. Figure S8 shows the distribution of correctly identified alignment edges. Within the TM-Vec held out dataset, the true positive distribution of proteins held out from the training roughly resembles the true positive distribution of proteins observed in pre-training. The average true positive rate, false positive rate and false discovery rate are shown in Table S3. As expected, TM-Vec performs best with the TM-align structural alignments on sequences that have been used for training the LSTM language model. This is observed in the true positive rate in addition to the false positive and false negative rates, as shown in Table S3. Thus, it appears that the generalization of TM-Vec primarily hinges on the underlying language model, as suggested by Rao et al [30].

### Embedding methods benchmarks

In Figure 3B, we compared TM-Vec to 6 other representations: one sequence-based, ProtTrans [32], and five different structure-based methods: cliques, GRAFENE [67], ORCA [68], CNN, influenced by DeepFRI [69], and GCN, influenced by the Kipf & Welling graph autoencoder (GAE) [70]. Each structure-based method in some manner consumes a thresholded distance matrix, or contact map, and is used to output a fixed-sized feature vector that is meant to encode structural information.

The three structure-based methods, cliques, GRAFENE, and ORCA, output so-called manually engineered features; in particular, these feature vectors are histograms over known non-redundant graph substructures called graphlets. We introduce cliques as a simple baseline, which consists of counting the ratio of non-overlapping cliques up to size 7 inside a given contact map. ORCA and GRAFENE count more advanced graphlet substructures including graphlet orbits (which consider the relative node identity within the graphlet).

We also evaluate against two other methods that admit learned, structure-based representations: Deep-FRI and the Kipf & Welling GAE. Each method consists of training an autoencoder on contact maps and extracting average-pooled representations from one of the hidden layers in the inference mode. DeepFRI is a CNN autoencoder whereas the GAE is a graph autoencoder. Both models are trained to minimize the binary cross entropy of the original contact map and its reconstruction.

Of the 5 selected structure-based methods, 4 of them are permutation invariant, with the exception being DeepFRI which considers the canonical sequence ordering and treats the input matrix as an image. Additionally, the manual crafted feature vectors do not scale well with graph density and hence cannot be evaluated for larger angstrom thresholds.

Evaluation metrics for Figure 3B include cluster adjusted mutual information and triplet scoring AUPR. Each benchmark is applied separately to the 4 CATH tiers to the top 5 most represented categories of that tier. In cluster adjusted mutual information, we apply spectral clustering using 5 clusters to the input feature vectors and calculate the adjusted mutual information between the cluster assignments and the actual label assignments. In triplet scoring AUPR, we choose triplets in which two of the three share the same label assignment while the third is drawn from a different category. We construct a balanced classification problem by considering the same-label pairs as the positive class and the same number of differently labeled pairs as the negative class. We use the cosine similarity among the selected positive and negative pairs as a classification prediction and calculate the AUPR.

## Software and Data Availability

TM-vec can be found at https://github.com/tymor22/tm-vec

DeepBLAST can be found at https://github.com/flatironinstitute/deepblast

## Acknowledgements

T.H. was funded by NIH R01DK103358, Simons Foundation, NSF-IOS-1546218, R35GM122515, NSF CBET-1728858, NIH R01AI130945. J.T.M was funded by the intramural research program of the Eunice Kennedy Shriver National Institute of Child Health and Human Development (NICHD). We would like to thank the Flatiron Institute, and particularly Ian Fisk, Nick Carriero, and Dylan Simon for providing the computer support required to train these models. We would like to thank Stephen Ra for helpful discussions. We thank the NIH, NSF, and Simons foundation for providing funding to pursue this work. We want to acknowledge Vikram Mulligan and Douglas Renfrew from the Flatiron Institute, Michael Heinzinger, Ahmedand Elnaggar, Christian Dallago, Konstantin Weibenow from TU Munich, Prathima Srinivas and Emily Webber from AWS, and Rob Knight and Igor Sfiligor from UCSD for their discussions. Lastly, we want to acknowledge Pytorch [71], Pytorch-Lightning [72], Biopython [73], Scipy [74], Numpy [75], and PyMOL [76] for providing software supporting scientific computing and visualization.

---

Table S1: Predicting TM-scores for novel folds discovered by the Microbiome Immunity project (MIP) There was a Pearson correlation of .786 between TM-Vec’s TM-score predictions and the known TM-scores for pairs of proteins from new, undiscovered folds discovered by the MIP project. This correlation is nearly identical to the correlation of 0.78 that we got predicting on a test dataset of left out folds from CATH.

Table S2: TM-Vec search accuracy on CATH datasets across multiple tiers with varying sequence similarity. Across every tier of CATH, and for two CATH datasets, CATH S40 (clusters thresholded at 40% sequence similarity) and CATH S100 (thresholded at 100% sequence similarity), we show the classification accuracy of the nearest neighbors returned by TM-Vec. For example, the Top 1 accuracy for Topology in CATH S100 is 97.7%, which means that 97.7% of the time, the nearest neighbor returned using TM-Vec is in the same fold as the query domain’s fold. As another example, the Top 3 column for Topology indicates the percentage of the time that one of the top 3 nearest neighbors returned by TM-Vec shares the same fold as the query domain’s fold.

Table S3: A breakdown of DeepBLAST’s performance on held out alignment datasets. Breakdown of the DeepBLAST True positive, False positive and False negative rates on the held out alignment datasets.

**Figure S1:**
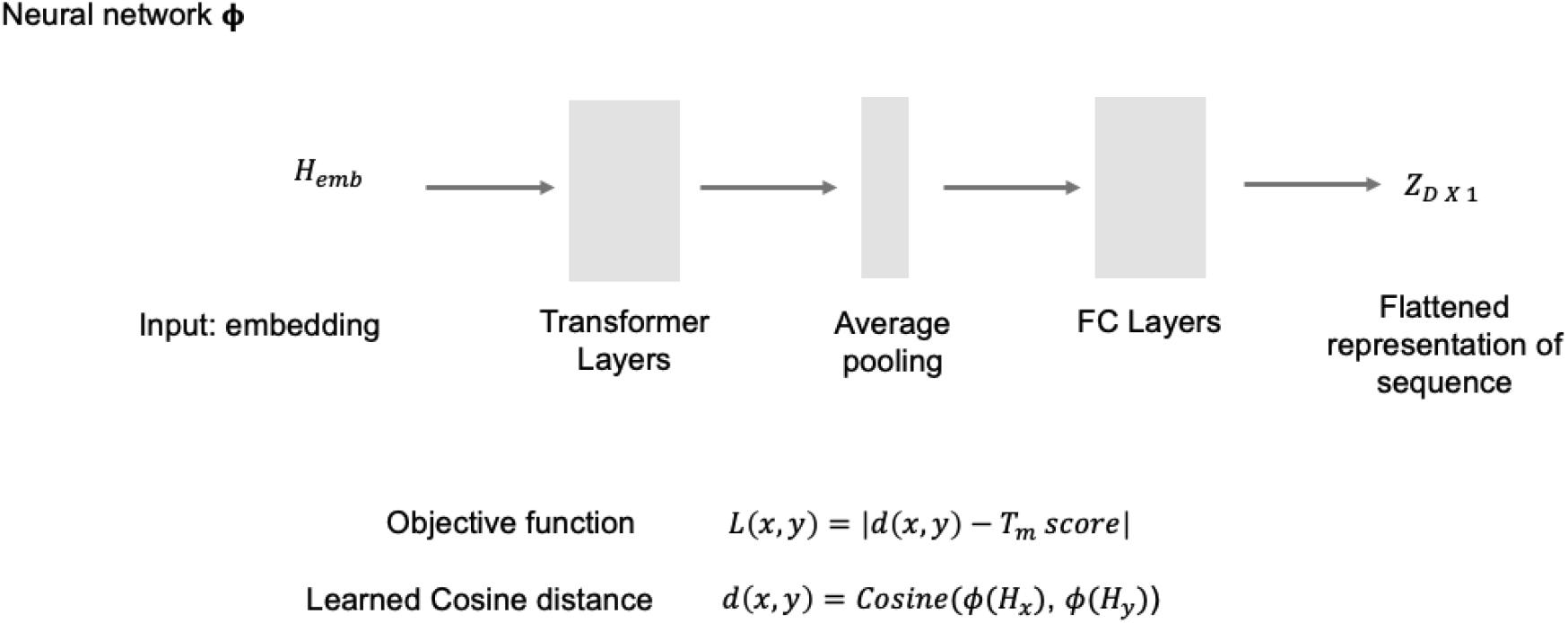
Overview of TM-Vec neural network architecture. The function ϕ takes in residue embeddings and produces a flattened vector representation for each protein. ϕ is composed of several transformer encoder layers, followed by average pooling, dropout, and fully connected layers. At the final step, we calculate the cosine distance between the vector representations of each protein in the pair, and our training objective is to minimize the L1 distance between the cosine similarity of the pairs vector representations, *z*, and their TM-score.

**Figure S2:**
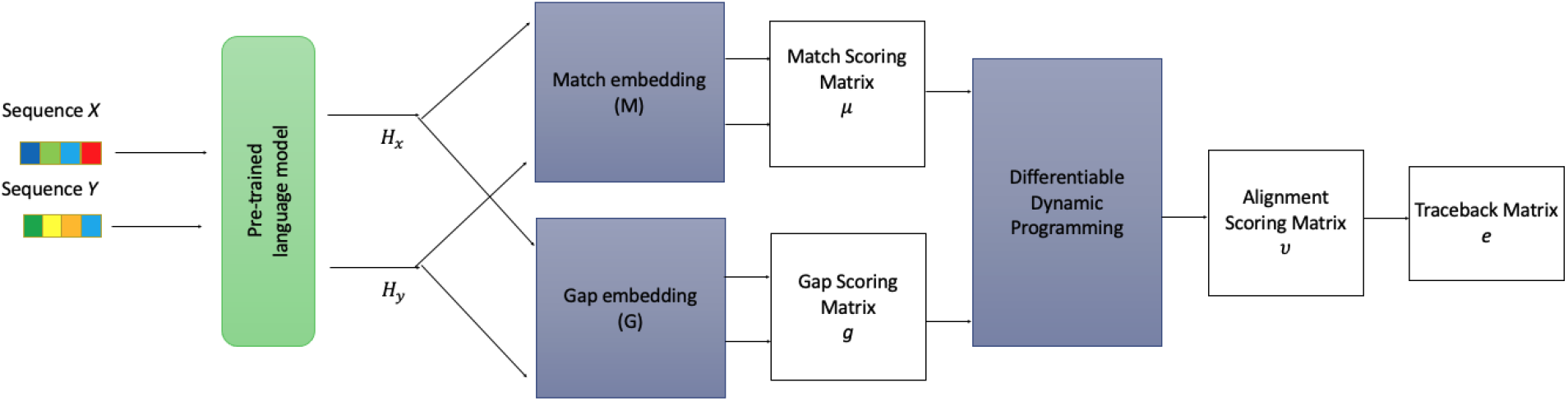
Overview of aligner pipeline. Proteins *X* and *Y* are fed into the pretrained LSTM protein language model [27] to obtain embeddings H_X_ and *H_Y_*. These residue-level embeddings are then propagated through the match embeddings (M) and gap embeddings (G) in order to obtain the match scores **μ** and the gap scores **g**. The match and gap scores are used to evaluate the differentiable dynamic programming algorithm and generate a predicted alignment traceback.

**Figure S3:**
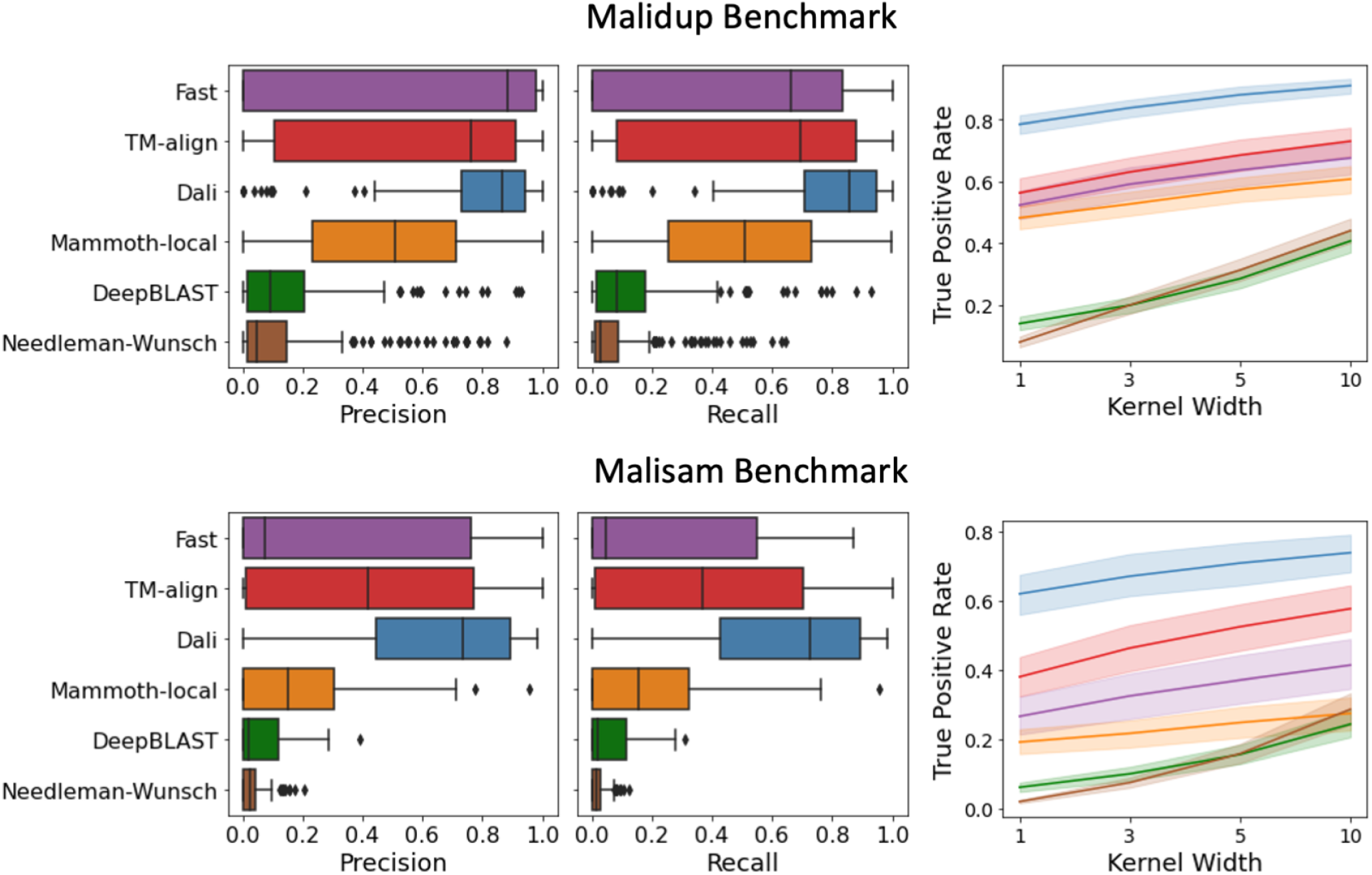
Precision and Recall metrics for each alignment on Malidup and Malisam benchmarks. The true positive rate was evaluated within window.

**Figure S4:**
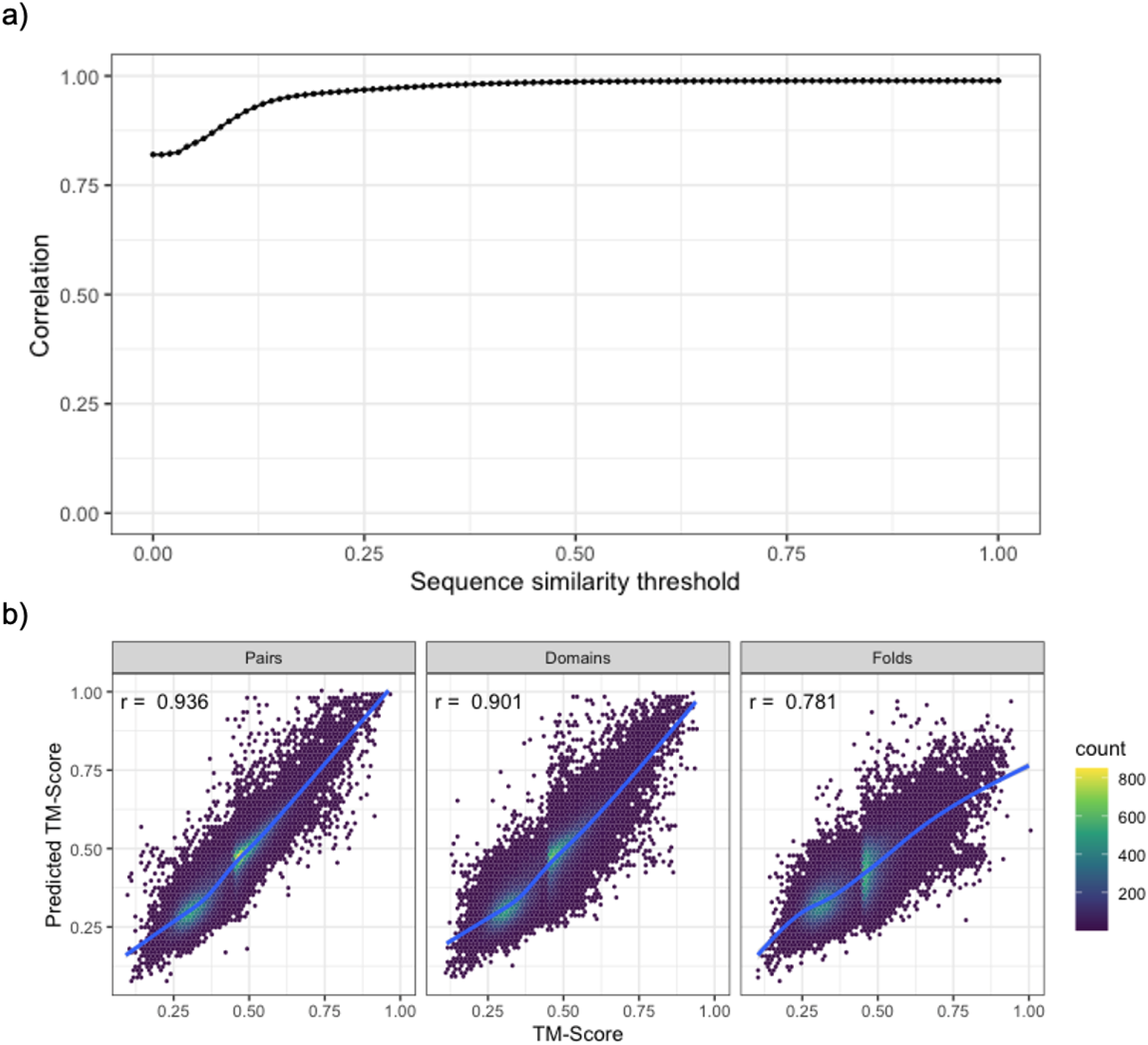
a) SwissProt cumulative Pearson correlations between the known TM-scores and predicted TM-scores for pairs of sequences at different sequence identity thresholds. The correlation coefficient at a particular value represents the correlation coefficient for SwissProt test pairs below that sequence identity threshold. b) CATH S40 predicted TM-scores versus ground truth TM-scores for different test datasets (Pairs, Domains, Folds). Contour plots show the density of points, and we also show the trend line for the relationship and Pearson correlation.

**Figure S5:**
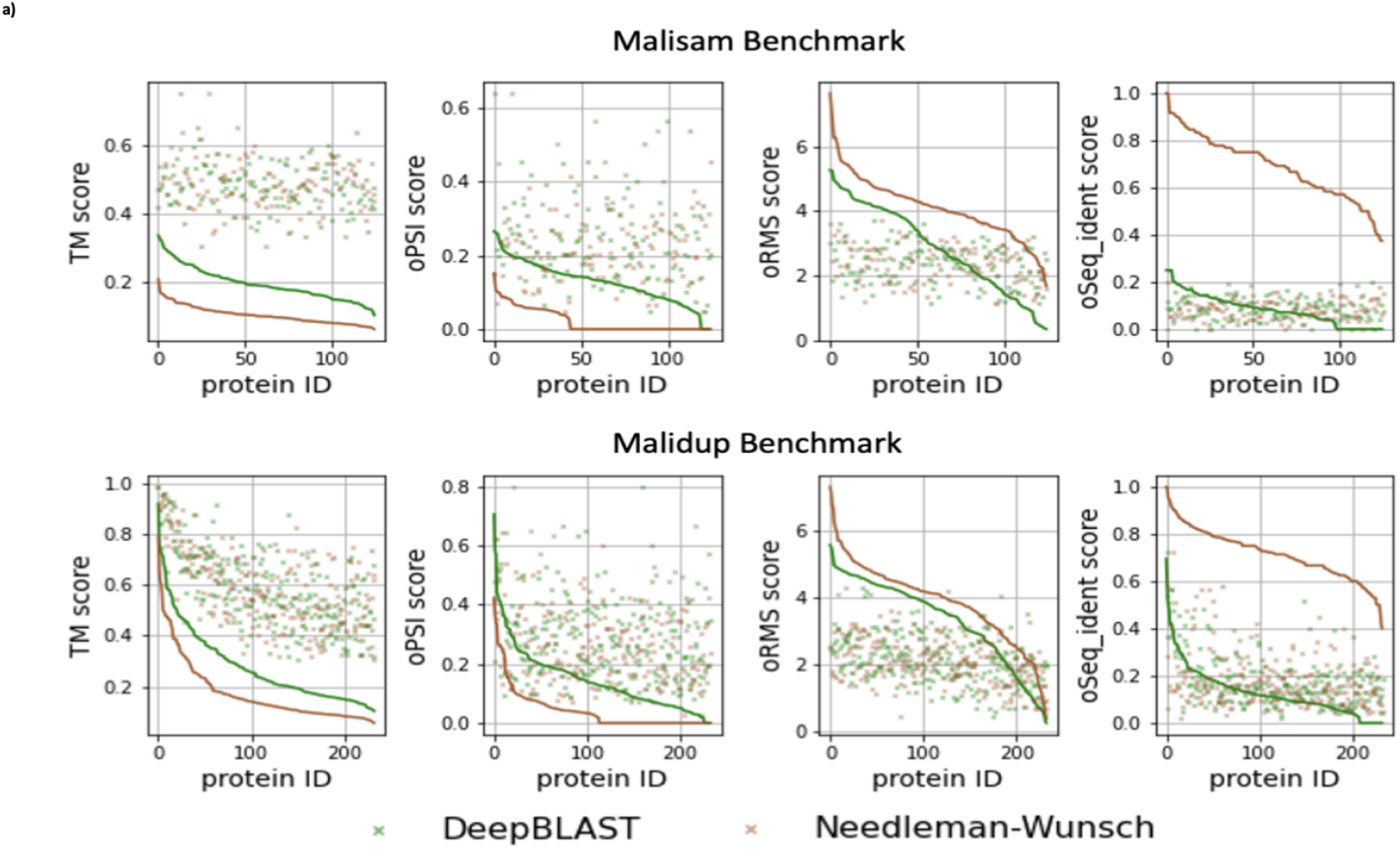
Comparison of DeepBLAST and Needleman-Wunsch on the Malidsam and Malidup benchmark. TM-score measures the superposition agreement between the two aligned protein structures. The oPSI metric measures the fraction aligned residues relative to the smaller protein on the aligned residues predicted to strongly superimposed by the alignment method. The oRMS metric measures the root mean standard deviation of the atomic positions on the aligned residues predicted to strongly superimposed by the alignment method. The oSeq identity score measures the fraction of identical sequence measured over the subset of the sequence alignment that was also aligned structurally by method. All of the alignment metrics are displayed in rank order, and the points represent the manual scores for that given protein, representing and upper or lower bound of the correct alignment.

**Figure S6:**
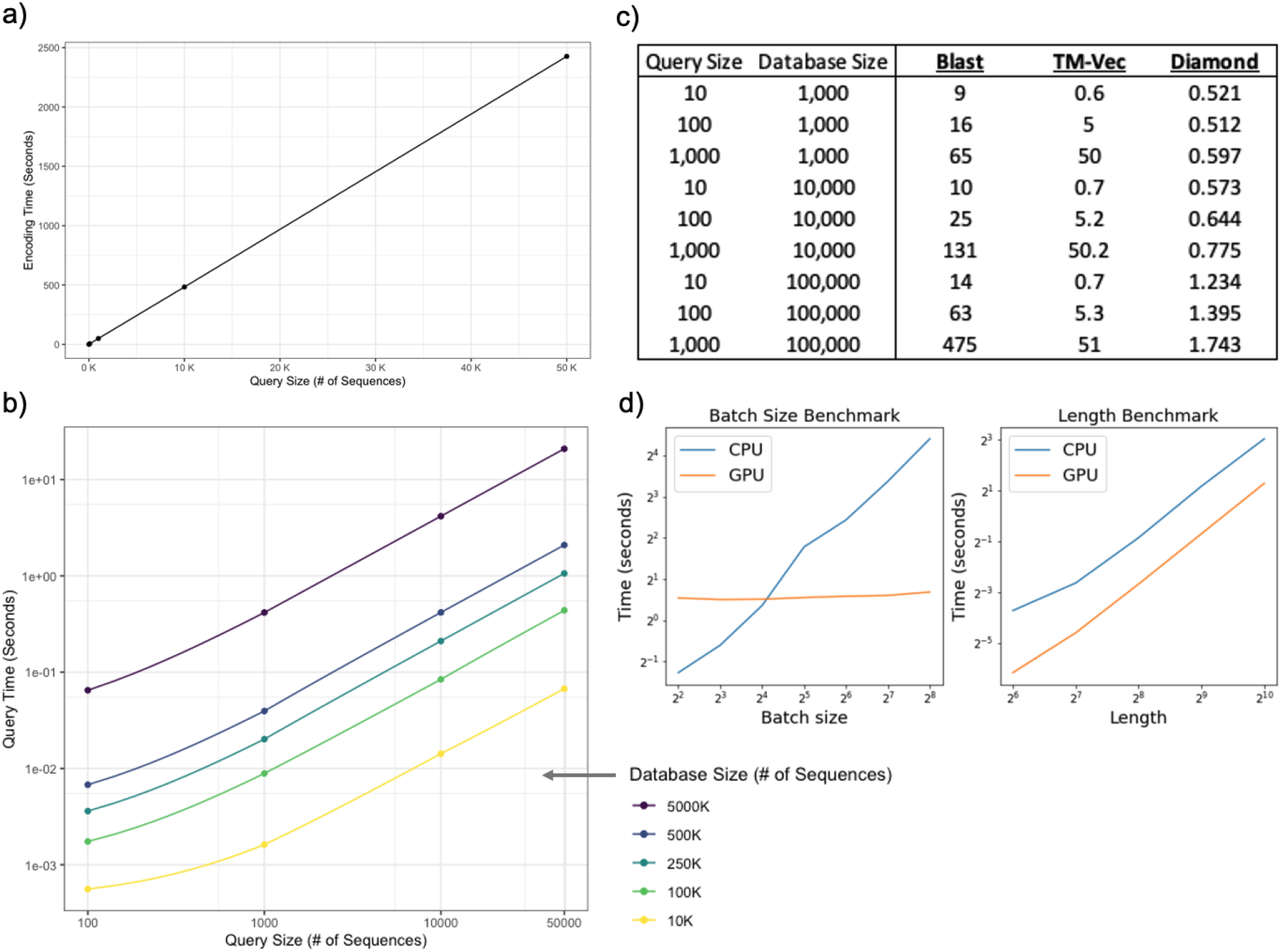
a) We evaluated how long it takes to encode query sequences using TM-Vec without parallelization on 1 GPU. The query size is on the x axis, and the encoding time in seconds is on the y axis. Encoding queries is linear in time. b) We evaluated how long it takes to search an indexed TM-Vec database and return the nearest neighbors once the queries have been encoded. The query size is on the x axis (number of sequences) and the query time is on the y axis (number of seconds) for databases of different sizes ranging from 10K sequences to 5M sequences. Search time is trivial relative to the time it takes to encode queries, and the size of the database does not materially impact speed. c) Search and alignment speed for TM-Vec is compared against BLAST, and Diamond for different query and database sizes. d) CPU vs GPU Differentiable Needleman-Wunsch benchmarks. The batch-size benchmark was run with randomized proteins of length 800 and the length benchmark was run with a fixed batch size of 64.

**Figure S7:**
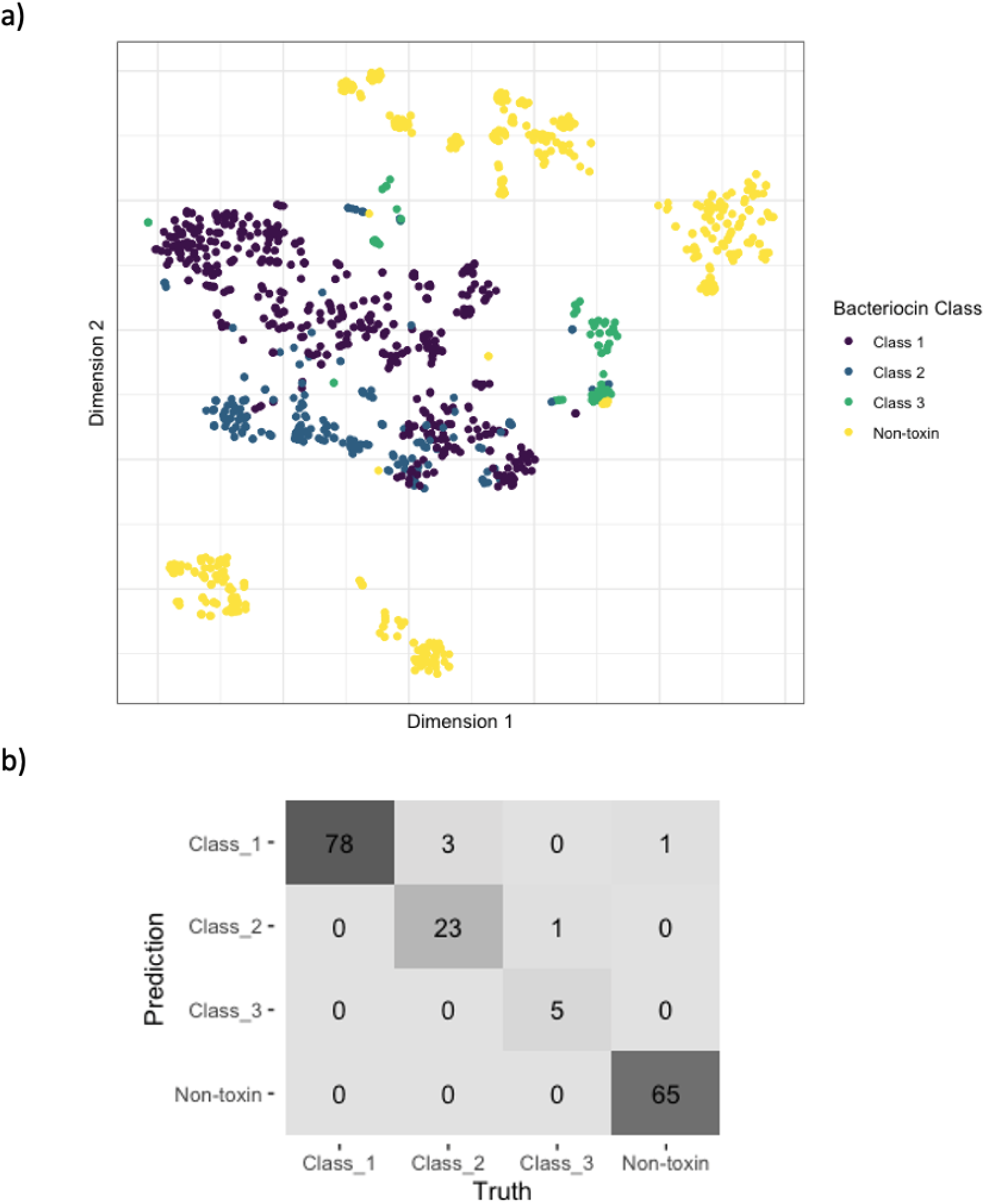
a) T-sne visualization of embeddings for both bacteriocins and non-toxin proteins. Clearly there is a separation of bacteriocins and non-bacteriocins. b) Confusion matrix for predicting the bacteriocin status/class for a held out test set of proteins. 85% of data was used to train a k-nearest neighbors classifier with k equal to 3 neighbors. The overall precision and recall on the held-out test set were 0.98 and 0.93, respectively.

**Figure S8:**
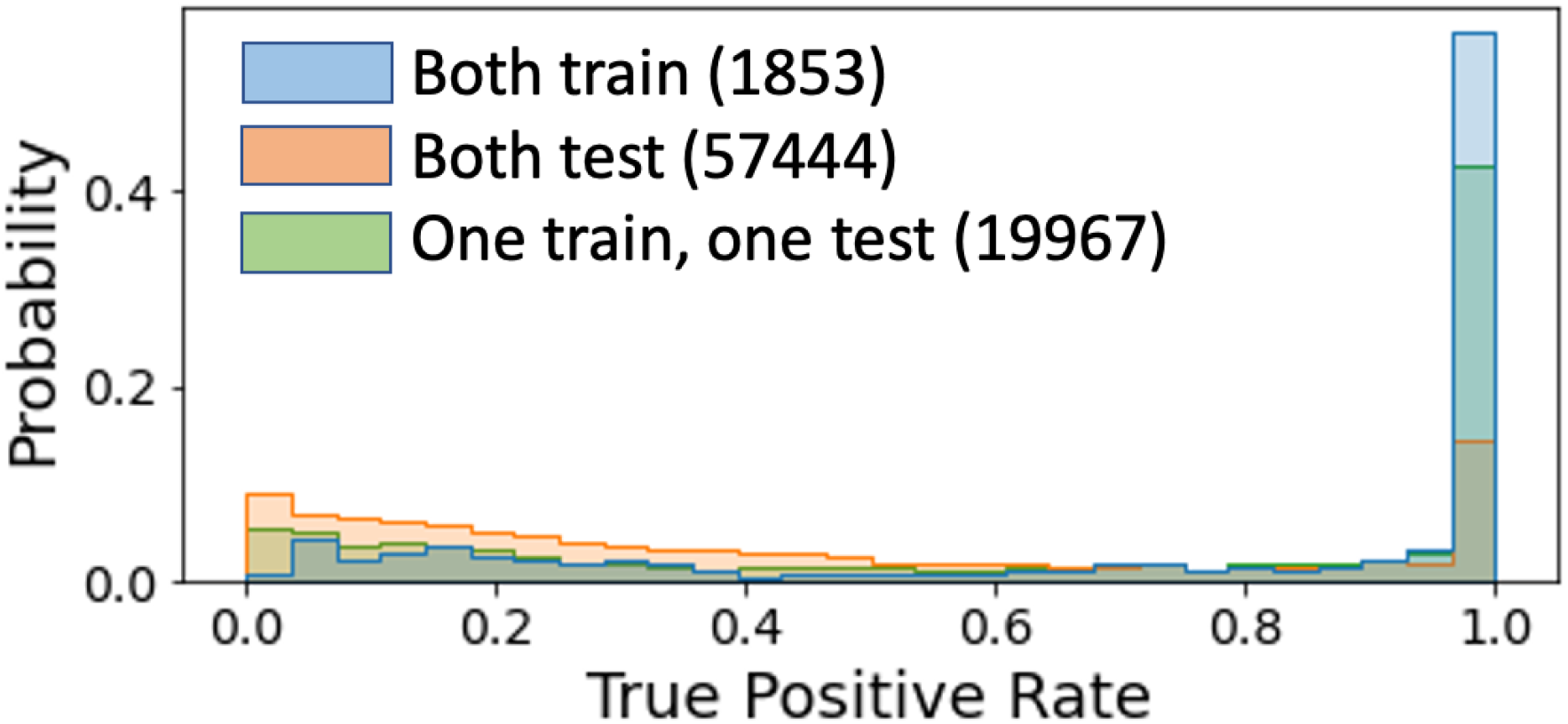
Distribution of TM-Vec true positive rates across heldout alignment datasets.

